# Programmable translational inhibition by a molecular glue-oligonucleotide conjugate

**DOI:** 10.1101/2025.07.15.664547

**Authors:** Siyi Wang, Joanna R. Kovalski, Francisco J. Zapatero-Belinchón, Maxwell Bennett, Duygu Kuzuoglu-Öztürk, Qiongyu Li, Erica Stevenson, Jie Liu, Nevan J. Krogan, Melanie Ott, Danielle L. Swaney, Davide Ruggero, Kevin Lou, Kevan M. Shokat

## Abstract

Selective inhibition of mRNA translation is a promising strategy for modulating the activity of disease-associated genes, yet achieving both high potency and specificity remains challenging. Rocaglamide A (RocA), a molecular glue, inhibits translation by clamping eIF4A onto polypurine motifs found in many transcripts, thereby limiting RocA’s specificity. Here, we developed RocASO, a chemical conjugate that links RocA to an antisense oligonucleotide (ASO) capable of base-pairing with defined mRNA sequences, thus directing RocA’s clamping mechanism to chosen targets and enhancing overall potency and specificity. We show that RocASOs are compatible with various types of ASO modalities, including gapmers that induce the degradation of target RNAs. RocASOs were designed to effectively knock down endogenous genes (*PTGES3*, *HSPA1B*) and SARS-CoV-2 viral RNA, the latter conferring potent antiviral activity in cells. These findings establish RocASO as a versatile platform for programmable translational inhibition with therapeutic potential.

## Main text

As the molecular blueprint for protein biosynthesis, RNA underlies numerous biological processes and is recognized as a high-value therapeutic target (*1–7*). Nonetheless, designing molecules that can bind RNA with high potency and specificity in the cellular environment remains challenging. Traditional small molecules typically target well-defined structural pockets, but these compounds can be difficult to apply to the dynamic, flexible nature of RNA and often fail to disrupt function through binding alone (*8, 9*). In contrast, antisense-based modalities, including small interfering RNAs (siRNAs) and antisense oligonucleotides (ASOs), leverage sequence complementarity for target recognition, although their clinical utility is often compromised by limited inhibitory potency, suboptimal intracellular delivery, and off-target effects such as unintended gene silencing or immune stimulation (*10–12*).

Despite these obstacles, ASOs have achieved notable clinical success, as evidenced by multiple FDA-approved therapies that employ various mechanisms, including RNase H-mediated target transcript degradation (*11, 13, 14*). For example, olezarsen, a clinically approved gapmer ASO, forms a partial DNA/RNA duplex upon binding its target transcript, which in turn is recognized by RNase H, leading to a reduction in hepatic apoC-III mRNA and corresponding triglyceride levels (*15*). However, the often relatively modest efficacy of ASOs has limited their broader therapeutic applications, particularly in contexts where robust target knockdown may be required. A broadly applicable, base-programmable strategy capable of binding and inhibiting RNA with enhanced potency would serve as a useful tool for modulating disease-associated RNAs, including exogenous RNAs such as viral RNA.

Here, we introduce RocASOs, a novel class of small molecule-oligonucleotide conjugates that chemically link ASOs to rocaglamide A (RocA). RocASOs combine sequence-specific RNA recognition via an ASO with molecular glue-mediated clamping of eIF4A to RNA via RocA (Fig. 1A). Unlike traditional small molecules, RocA binds the interface formed by eIF4A and RNA, inhibiting translation by blocking ribosomal scanning in the 5′ untranslated region (UTR) (*16–18*). However, despite RocA’s preference for polypurine motifs (i.e., purine-rich sequences), these motifs are broadly distributed across transcripts, leading to widespread translational inhibition and dose-limiting toxicity. By conjugating RocA to an ASO designed to bind sequences adjacent to a polypurine motif, the resulting bivalent RocASO can direct the eIF4A clamp to a specific transcript, potentially enhancing overall binding affinity and target selectivity.

**Fig. 1.**
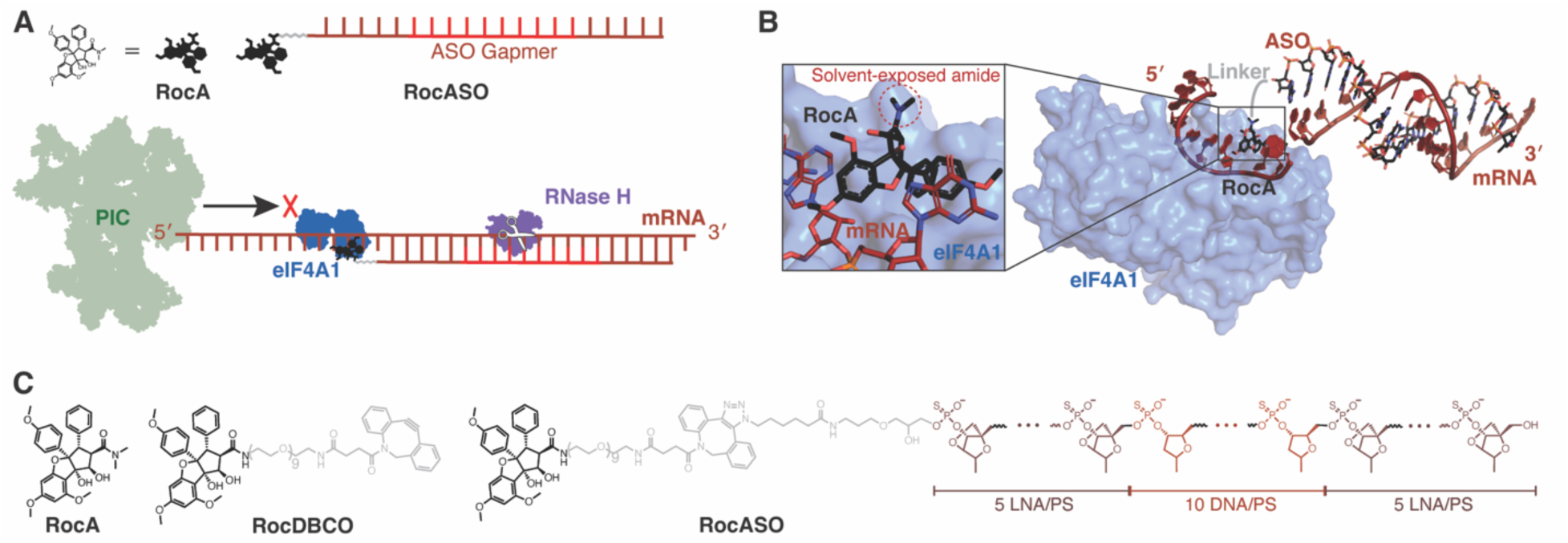
RocASO design and proposed mechanism of action. **(A)** Schematic representation of RocASO’s design and proposed mechanism of action. The ASO component directs RocA to specific transcripts, increasing local RocA concentration and improving blocking of the pre-initiation complex (PIC), while RocA stabilizes the ASO-mRNA duplex by clamping eIF4A onto a polypurine motif, preventing ASO displacement and enhancing RNA degradation. **(B)** Structural model of the eIF4A1-RocASO-mRNA complex (PDB: 5ZC9 and 4KYY combined), illustrating the molecular interactions responsible for translation inhibition. A zoomed-in view highlights the solvent-exposed C2 amide of RocA. **(C)** Chemical structures of RocA, RocDBCO, and RocASO. RocA core (black), linker (gray), DNA nucleotide segments (red), and locked nucleic acid (LNA) nucleotide segments (dark red) of RocASO’s structure are depicted.

### Design and synthesis of RocASOs

Guided by structural insights from the eIF4A1-RocA-RNA ternary complex (*17*), we identified a suitable modification site on RocA for conjugation to an ASO. The crystal structure revealed that RocA’s C2 *N*,*N*-dimethylamide group is fully solvent-exposed, making it an ideal site for conjugation since chemical modifications at this position would not be expected to disrupt any binding interfaces (Fig. 1B). Supporting this hypothesis, prior studies demonstrated that chemical substitution at this site has minimal impact on RocA’s activity (*19–22*). Thus, we aimed to generate a bivalent molecule by attaching RocA to a sequence-specific ASO through a modified C2 amide.

To achieve this, we synthesized RocDBCO, a derivative of RocA functionalized with a dibenzocyclooctyne (DBCO) group at the solvent-exposed amide via a flexible polyethylene glycol (PEG) linker (Fig. 1C). A PEG9 linker was chosen for sufficient flexibility and length to enable simultaneous binding of the RocA-eIF4A-mRNA complex and the ASO-mRNA duplex in tandem (Fig. 1B). Although RocDBCO exhibited slightly reduced potency – likely due to altered cellular uptake or target engagement – it maintained cellular permeability and dose-dependent inhibition of RocA-sensitive transcripts (fig. S1A, B). These results confirmed that introducing a modification at the C2 amide of RocA preserves its functional activity, validating RocDBCO as a suitable scaffold for subsequent ASO conjugation.

Finally, RocDBCO was reacted with azide-modified ASOs (locked nucleic acid (LNA) gapmer, phosphorothioate linkage) via strain-promoted azide-alkyne cycloaddition (SPAAC) (*23*) to furnish the desired RocASOs (Fig. 1C, fig. S1C, D). In this design, the ASO hybridizes adjacent to a polypurine motif, localizing RocA’s activity by increasing its effective concentration near a specific clampable sequence. Moreover, this configuration may also stabilize the ASO-mRNA duplex through the formation of a RocA-eIF4A-RNA ternary complex directly upstream, potentially preventing ASO displacement and enhancing overall translational inhibition (Fig. 1A).

### Evaluation of RocASOs targeting *PTGES3*

To evaluate the activity of RocASO, we selected prostaglandin E synthase 3 (*PTGES3*), a cochaperone which also plays a role in prostaglandin E2 biosynthesis, as a representative target gene (*24, 25*). The *PTGES3* mRNA contains multiple polypurine motifs in its 5′ UTR (Fig. 2A) and is among the most differentially translated transcripts following RocA treatment (*16*), indicating its suitability as a substrate for eIF4A clamping. Endogenous PTGES3 protein levels were directly monitored using a split-mNeonGreen fluorescent reporter integrated into the native *PTGES3* locus (*26*). We first assessed the effects of RocA and its derivative RocDBCO on PTGES3 expression. RocA decreased PTGES3 protein levels at low concentrations but exhibited a mild rebound effect at higher doses, potentially due to an adaptive response to cell stress from non-specific translation suppression (*27*). In contrast, RocDBCO – the direct precursor of RocASO – did not inhibit PTGES3 reporter levels at the concentrations tested, consistent with its reduced potency observed in western blot and cell viability assays (Fig. 2B, fig. S1A, B).

**Fig. 2.**
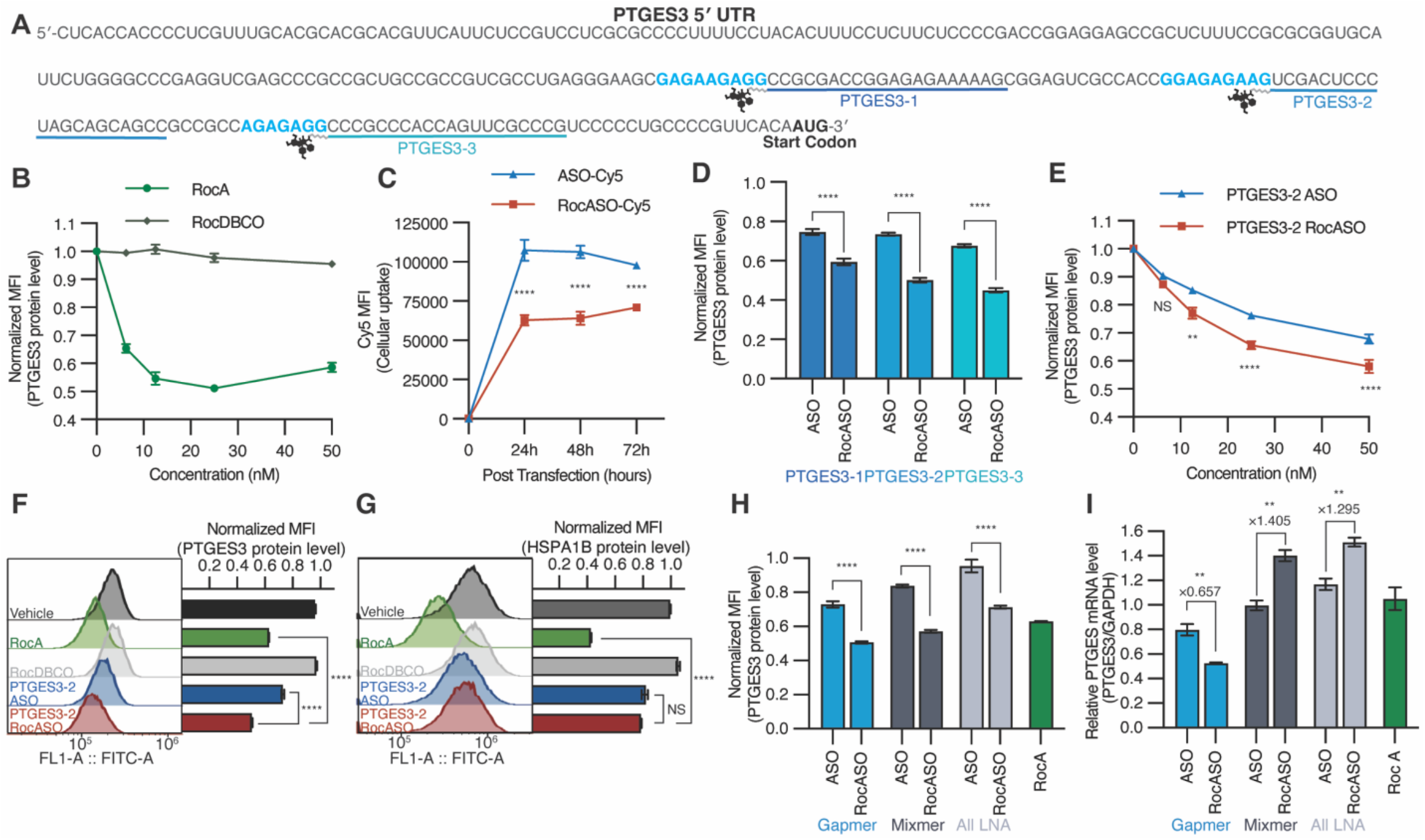
Design and functional characterization of *PTGES3*-targeting RocASOs. **(A)** *PTGES3* mRNA 5′ UTR sequence highlighting polypurine motifs (blue) and RocASO target sites (underlined). **(B)** PTGES3 protein expression as measured by fluorescence flow cytometry after 48 hours of the indicated treatments in HEK293T split-mNeonGreen (mNG) cells, where endogenous PTGES3 has been tagged. **(C)** Transfection efficiency of PTGES3-1 ASO and RocASO (10 nM), measured by Cy5 fluorescence intensity via flow cytometry at indicated time points. ASOs were labeled with Cy5 at the 5′ end. **(D)** Normalized fluorescently tagged PTGES3 protein levels after indicated treatment (10 nM, 48 hours post-transfection). **(E)** Dose-dependent effects of PTGES3-2 ASO and RocASO on PTGES3 protein levels (48 hours post-transfection). **(F, G)** Fluorescent spectra and normalized PTGES3 **(F)** and HSPA1B **(G)** fluorescently tagged protein levels after indicated treatment (10 nM, 48 hours post-transfection). **(H, I)** Effects of different ASO designs and their corresponding RocASOs on PTGES3 protein (H) and mRNA (I) levels (10 nM, 48 hours post-transfection). Unless stated otherwise, data are presented as mean ± SEM (n = 3 biological replicates), and statistical analysis was performed using two-way ANOVA. All treatments were normalized to vehicle control (transfection reagent + 0.1% DMSO). Flow cytometry data are represented by median fluorescence intensity (MFI).

Next, we sought to design RocASOs targeting the *PTGES3* mRNA by identifying polypurine motifs within its 5′ UTR. In this initial study, polypurine motifs were defined as sequences of six or more consecutive purine bases, a pattern previously demonstrated to be sufficient for RocA to clamp eIF4A onto RNA (*16*). We identified three such motifs in the *PTGES3* 5′ UTR and synthesized three corresponding RocASOs (PTGES3-1, -2, -3), each employing an LNA gapmer ASO format and designed to bind sequences immediately downstream of a polypurine motif (Fig. 2A). To assess the impact of RocA conjugation on cellular uptake, we also synthesized a RocASO derivative labeled with Cy5 and compared its uptake to that of the corresponding ASO-Cy5. RocASO-Cy5 treated cells displayed robust Cy5 fluorescence, albeit lower than that in ASO-Cy5 treated cells (Fig. 2C). This suggested that RocASOs retain the ability to be transfected into cells, though with modestly reduced intracellular concentrations compared to ASOs.

Despite this reduced cellular uptake, all three RocASOs significantly enhanced PTGES3 protein knockdown compared to their corresponding unconjugated ASOs, with PTGES3-2 and PTGES3-3 RocASOs showing the most substantial improvements (Fig. 2D, fig. S2A). Dose-dependent analysis further corroborated these results, demonstrating that all three RocASOs generally outperformed their ASO counterparts across various concentrations (Fig. 2E, fig. S2B, C). We next measured *PTGES3* transcript levels by qPCR following treatment with the most potent RocASOs, PTGES3-2 and PTGES3-3, and observed a greater reduction in RNA levels compared to their unconjugated gapmer ASO counterparts. These results suggest that enhanced RNA degradation contributed to the improved knockdown efficacy of RocASOs (fig. S2D).

### Mechanistic characterization of RocASO activity and specificity

We systematically modified various components of our RocASO design to determine their contributions to overall activity. To evaluate the role of ASO positioning relative to the polypurine motif, we engineered RocASOs in which the ASO binding site was shifted upstream of the polypurine motif (PTGES3-4 and PTGES3-5). In these variants, RocA was conjugated to the 5′ end rather than the 3′ end of the ASO to ensure that RocA remained in close proximity to the polypurine motif (fig. S3A). However, neither of these examples enhanced mRNA degradation or protein suppression (fig. S3B, C, D). Similarly, a RocASO targeting a sequence 10 bases distal from the closest polypurine motif (PTGES3-6) also did not demonstrate improved RNA degradation or protein level inhibition (fig. S3B, C, D). Collectively, these findings indicate that optimal translational inhibition requires the RocA binding site to reside within a linker-accessible distance upstream of an ASO-complementary tract. We propose that this configuration enables RocA to effectively engage in tandem with the oligonucleotide–RNA duplex, facilitating the formation of a stable RocA–eIF4A–polypurine mRNA complex upstream of the ASO–RNA duplex that prevents displacement by scanning helicases or ribosomes (fig. S3E).

RocA broadly suppresses translation by targeting short polypurine motifs found in many transcripts, resulting in limited specificity and dose-limiting toxicity. To evaluate whether RocASOs improve target selectivity, we examined their effects on a potential off-target gene, heat shock protein family A (Hsp70) member 1B (*HSPA1B*), using a split-mNeonGreen fluorescent reporter similar to that for *PTGES3*. This *HSPA1B* mRNA also contains multiple polypurine motifs in its 5′ UTR and it has similarly been shown to be susceptible to RocA-mediated clamping (*16*). Treatment with a *PTGES3*-targeting RocASO had minimal impact on HSPA1B protein levels, comparable to its corresponding unconjugated ASO (Fig. 2F, G). In contrast, RocA markedly suppressed HSPA1B protein expression (Fig. 2G), indicating that RocASOs achieve improved selectivity through sequence-specific base-pairing mediated by their conjugated complementary oligonucleotides.

Lastly, we confirmed that RocASO-induced RNA degradation operates through a gapmer ASO mechanism, which involves partial DNA/RNA hybridization and RNase H-mediated cleavage, by designing two additional PTGES3-2 RocASOs in mixmer (i.e., alternating LNA and DNA bases) and all-LNA formats. All RocASOs, regardless of ASO format, enhanced protein knockdown compared to their corresponding ASOs (Fig. 2H). However, only the gapmer ASO and its RocASO counterpart reduced *PTGES3* mRNA levels, with RocASO demonstrating greater transcript and protein knockdown. In contrast, the mixmer and all-LNA ASOs, which inhibit translation via steric blocking, did not alter *PTGES3* mRNA levels, and their corresponding RocASOs even increased mRNA levels, which may reflect compensatory transcriptional upregulation (Fig. 2I). Despite this increase in target mRNA levels, steric-blocking RocASOs still exhibited more robust protein knockdown than their ASO counterparts. Altogether, these data suggest that RocA conjugation may enhance ASO-mediated knockdown by increasing steric blocking efficiency and/or enhancing RNA degradation through stabilizing the mRNA-bound complex.

### Reprogramming RocASOs to target *HSPA1B*

To investigate whether the RocASO platform can be generalized to additional transcripts, we designed four RocASOs targeting the 5′ UTR of *HSPA1B* mRNA, each directed at a distinct polypurine motif. As with our most potent RocASOs tested thus far, RocA was conjugated to the 3′ end of each ASO to ensure optimal positioning (fig. S4A). In our endogenously tagged fluorescent reporter assay, none of the unconjugated ASOs measurably reduced HSPA1B protein levels, whereas all corresponding RocASOs significantly suppressed HSPA1B protein expression; a scrambled control RocASO had no effect (fig. S4B). These findings demonstrate that RocASO-mediated translational suppression is generalizable and can be applied to different targets when designed according to the positioning principles established in the *PTGES3*-targeting study.

### Design and validation of a RocASO targeting SARS-CoV-2

Having demonstrated RocASO activity against endogenous human transcripts, we next sought to expand the target scope to exogenous viral mRNAs. Viruses rely on host machinery for translation and replication, often hijacking ribosomes and specific translation factors to preferentially engage viral transcripts (*28*). For example, a SARS-CoV-2 protein interaction map identified Nsp9 as an interactor of eIF4H, a cofactor that enhances eIF4A helicase activity – an essential component of viral translation (*29*). Viruses are thought to exploit eIF4A to facilitate translation of their mRNAs, particularly those with highly structured 5′ UTRs (*28*). Notably, RocA and its analogs have been shown to inhibit a range of viruses, including SARS-CoV-2 (*30*), suggesting that viral transcripts may be susceptible to the eIF4A clamping mechanism. Based on these observations, we hypothesized that RocA could engage polypurine motifs within the SARS-CoV-2 5′ UTR to suppress viral translation, and that incorporating ASO-mediated sequence recognition, as in our RocASO platform, would confer enhanced potency and selectivity for viral over host translation.

Candidate RocA-binding sites within SARS-CoV-2 mRNA were identified by searching for 6-mer motifs characterized as RocA binding sequences from previous RNA interaction studies (*16*). Guided by the design principles established for endogenous human transcripts, we designed a RocASO targeting one such polypurine motif (i.e., RocA binding sequence) within the SARS-CoV-2 5′ UTR (Fig. 3A). Sequence alignment across viral strains confirmed that both the polypurine motif and the adjacent ASO binding site are highly conserved, supporting the potential for broad activity of this RocASO against multiple SARS-CoV-2 variants (fig. S5A).

**Fig. 3.**
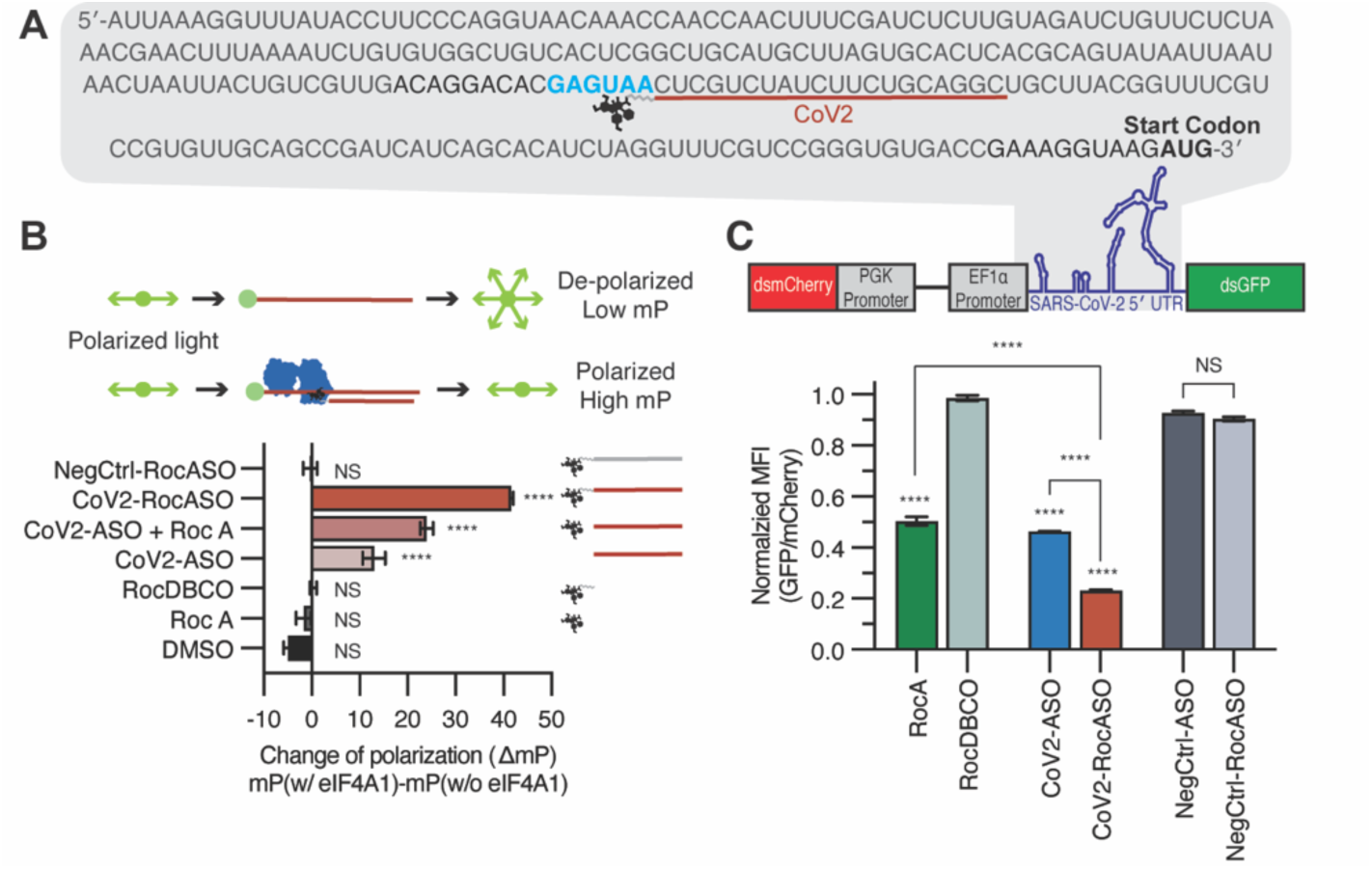
A SARS-CoV-2-targeting RocASO promotes sequence-directed ternary complex formation. **(A)** SARS-CoV-2 5′ UTR sequence highlighting polypurine motifs (blue) and RocASO target sites (underlined). **(B)** Fluorescence polarization (FP) assay illustration and results. Change of polarization values (ΔmP) indicates FP change in the presence or absence of 500 nM eIF4A1 protein with 1 µM of the indicated treatment and 10 mM AMP-PNP. **(C)** Illustration and results of the dual-reporter assay system used to measure SARS-CoV-2 5’ UTR–driven translation. dsmCherry: destabilized mCherry; dsGFP: destabilized GFP. Results are normalized by the GFP/mCherry ratio to depict specific effects on SARS-CoV-2 5’ UTR–driven translation (50 nM, 16 hours post-transfection). Unless stated otherwise, data are presented as mean ± SEM (n = 3 biological replicates), and statistical analysis was performed using two-way ANOVA. All treatments are normalized to vehicle control (transfection reagent + 0.1% DMSO). Flow cytometry data are represented by median fluorescence intensity (MFI).

To assess whether RocA or a RocASO can facilitate eIF4A1 recruitment to a SARS-CoV-2 RNA sequence, we performed a fluorescence polarization (FP) assay with an RNA probe derived from a segment of the SARS-CoV-2 5′ UTR that includes both the target polypurine motif and the adjacent ASO binding site. In this biochemical assay, an increase in FP in the presence of eIF4A1 indicates complex formation with the RNA. At the concentrations tested, neither RocA alone nor RocDBCO enhanced eIF4A1-RNA binding, potentially due to the longer probe length and/or lower ligand concentrations used here compared to previous studies (*16*) (Fig. 3B). A DNA-based ASO, either alone or when combined with free RocA, produced modest increases in FP signal, whereas the DNA-based CoV2-RocASO elicited the largest signal increase (Fig. 3B). These results suggest that RocA and ASO components can act cooperatively, and when linked together in a RocASO, form a stable, bivalent ternary complex – providing a biophysical rationale for the enhanced translational inhibition observed in cellular assays. Additionally, this binding was sequence-specific, as a non-complementary NegCtrl-RocASO failed to show binding (Fig. 3B). We further corroborated these findings using an LNA-modified gapmer RocASO - the modality used in most of our cellular assays, which exhibited comparable activity to the DNA-based construct (fig. S5B).

Next, we developed a dual-reporter HEK293T system to evaluate RocASO’s impact on SARS-CoV-2 5′ UTR–driven translation in cells. In this assay, destabilized GFP is expressed under control of the SARS-CoV-2 5’ UTR, while a destabilized mCherry, driven by a minimal 5′ UTR, functions as an internal control for global translation. Treatment with CoV2-RocASO significantly decreased the GFP/mCherry ratio compared to the unconjugated ASO, whereas a scrambled RocASO was inactive – findings that mirror the affinity and specificity observed in our biochemical experiments (Fig. 3C). This enhanced efficacy was again validated in dose-response, suggesting that the viral 5′ UTR might be similarly susceptible to translation inhibition by RocASOs (fig. S5C).

### Antiviral efficacy of a RocASO in a cellular model of SARS-CoV-2 infection

CoV2-RocASO’s antiviral efficacy and selectivity were evaluated using a live cell-based imaging assay in replication-competent SARS-CoV-2 infected cells treated with RocA, CoV2-ASO, or CoV2-RocASO (Fig. 4A). Consistent with previous reports, RocA inhibited viral infection in a dose-dependent manner; however, this effect was accompanied by a marked reduction in cellular confluency, suggesting a narrow therapeutic window between infection suppression and inhibition of global cellular translation (fig. S6A). In contrast, CoV2-ASO showed minimal impact on SARS-CoV-2 replication across the range of concentrations tested while remaining largely non-toxic (fig. S6B). Notably, CoV2-RocASO inhibited viral replication in a dose-dependent manner while also preserving cell viability – at 40 nM, infection was reduced to levels comparable to those achieved by RocA, yet cell viability remained comparable to that observed with CoV2-ASO (fig. S6C). Furthermore, normalizing SARS-CoV-2 infection to cellular confluency revealed that only CoV2-RocASO exhibited virus-specific inhibition, suggesting that the molecular glue-oligonucleotide conjugate selectively disrupts SARS-CoV-2 replication and translation (Fig. 4B).

**Fig. 4.**
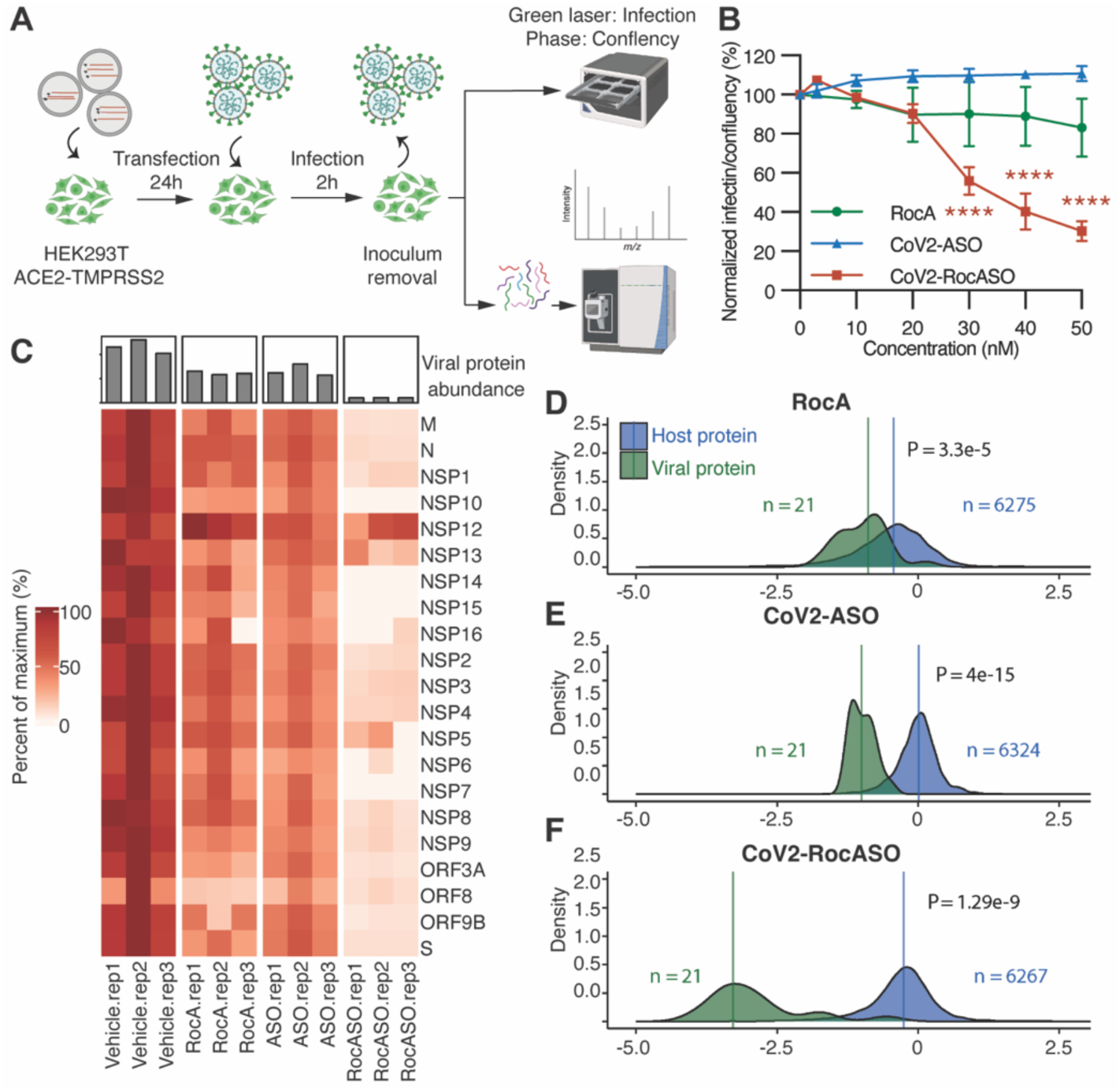
Antiviral efficacy of a RocASO in SARS-CoV-2–infected cells. **(A)** Experimental schematic. HEK293T-ACE2-TMPRSS2 cells were given the indicated treatments one day prior to SARS-CoV-2 infection. Infection and confluency were monitored using the Incucyte system, and in a separate experiment cells were harvested for proteomics analysis one day post-infection. **(B)** Specific antiviral effect of the indicated treatments, normalized by vehicle control (transfection reagent + 0.1% DMSO). Data are presented as mean ± SEM (n = 4 biological replicates), and statistical analysis was performed using two-way ANOVA. **(C)** Heatmap of viral protein abundances following the indicated treatments (40 nM), where each row represents a viral protein. Values are colored based on the percentage of the maximum observed abundance for that protein. The top bar chart indicates the mean viral protein abundance for each treatment condition. **(D–F)** Distribution of protein fold changes under RocA **(D)**, CoV2-ASO **(E)**, and CoV2-RocASO **(F)** treatment relative to vehicle control. Fold changes of host (blue) and viral (green) proteins are plotted; vertical lines indicate median values. Median log₂ fold changes were as follows: RocA (host: –0.43, viral: –0.89), CoV2-ASO (host: 0.01, viral: –1.00), and CoV2-RocASO (host: –0.26, viral: –3.28). Statistical comparisons were made using two-sided t-tests (n = 3 biological replicates).

To test this hypothesis, we performed quantitative proteomic analysis on SARS-CoV-2 infected cells treated with 40 nM of each agent. First, we examined each treatment’s ability to suppress viral proteins. Both RocA and CoV2-ASO treatment resulted in moderate reductions in viral proteins; however, treatment with CoV2-RocASO nearly abolished viral protein expression (Fig. 4C). Next, we evaluated the treatment impacts on global translation by quantifying the relative abundance of over 6000 host proteins (Fig. 4D–F, fig. S7A–C). In contrast to RocA, which broadly reduced host protein levels, and CoV2-ASO, which had virtually no effect, CoV2-RocASO mildly altered host protein expression while resulting in the greatest separation between viral and host protein effects. This result underscores the increased specificity of CoV2-RocASO relative to RocA and its enhanced potency relative to CoV2-ASO. Lastly, to assess the contribution of viral infection on these host protein changes, we conducted parallel quantitative proteomic analysis on uninfected cells under the same treatment conditions. In both RocA and CoV2-RocASO treated uninfected cells, host protein synthesis was slightly less affected compared to infected cells; however, CoV2-RocASO consistently exhibited a milder effect on host protein expression than RocA, even in the absence of infection (fig. S7D). These results support CoV2-RocASO’s potential as a selective antiviral agent with a favorable therapeutic window.

## Discussion

In this study, we developed RocASO, a programmable molecular glue-oligonucleotide conjugate that inhibits mRNA translation with high potency and specificity. RocASO combines RocA, a molecular glue that clamps eIF4A onto polypurine motifs, with a sequence-specific ASO, integrating multiple modes of mRNA inhibition into a single molecule. This design creates a cooperative interplay between the two components: (i) the ASO directs RocA’s molecular glue activity to selected sequences, limiting off-target engagement and improving specificity, and (ii) RocA stabilizes the overall RocASO-RNA assembly by forming a RocA-eIF4A-RNA ternary complex directly adjacent to the ASO-RNA duplex, which can enhance overall translational inhibition and transcript degradation. Furthermore, we establish the versatility and programmability of RocASOs to target a diverse set of mRNAs by recognizing polypurine motifs and hybridization sequences in tandem, and demonstrate their therapeutic potential in a cellular model.

By targeting exogenous viral RNA, RocASOs can effectively block viral translation, an approach that may be especially valuable against SARS-CoV-2. Although several therapies exist to address acute COVID-19, there are no approved treatments for long COVID, which continues to affect millions of people worldwide. Although the underlying mechanisms remain incompletely understood, accumulating evidence suggests that a persistent SARS-CoV-2 reservoir may contribute to prolonged symptoms in many patients (*31–33*). Lingering viral RNA can be actively translated, triggering ongoing immune activation even when widespread replication has ceased. By suppressing its translation and facilitating viral RNA degradation, RocASOs could represent a promising therapeutic strategy for the treatment of long COVID. Moreover, given the established delivery systems and the programmable nature of ASOs, this platform could be readily adapted to other RNA virus infections—particularly hepatotropic viruses—and may prove valuable in future emerging pandemics.

Despite exhibiting lower cellular uptake than unconjugated ASOs under our transfection conditions, optimally designed RocASOs consistently achieved enhanced target knockdown, suggesting they may require lower intracellular concentrations for therapeutic efficacy. Delivery-enhancement strategies commonly used for oligonucleotide therapeutics, such as GalNAc conjugation (*34, 35*) or lipid nanoparticle encapsulation, could further promote RocASO uptake in target tissues(*36*). Additionally, chemical optimization of RocA, the linker, and the ASO components may improve RocASO stability, potency, and biodistribution. Clinically validated ASOs like mipomersen and inotersen, which are intrinsically deliverable and hybridize to target sequences adjacent to short purine-rich tracts, present a promising scaffold to directly apply the RocASO strategy to boost therapeutic efficacy. Moreover, because RocASOs leverage dual recognition via RocA and ASO components, they may be compatible with shorter oligonucleotide sequences than those typically used in standard ASO modalities, potentially enabling more compact designs for broader therapeutic applications.

## Acknowledgements

We thank the Manuel D. Leonetti laboratory for generously providing the endogenously tagged fluorescent cell lines. We are grateful to the Jennifer Doudna laboratory— especially Abdullah Syed—for sharing their SARS-CoV-2 reporter replicon for initial testing and for providing the HEK293T-ACE2-TMPRSS2 cell line. We also thank Taha Y. Taha for the generous gift of the SARS-CoV-2 reporter replicon.

## Author contributions

S.W., K.L., and K.M.S. conceived the project, designed experiments, interpreted results, and wrote the manuscript. S.W. performed the chemical synthesis, RocASO design and conjugation, flow cytometry assays, qRT-PCR experiments, fluorescence polarization assays, immunoblotting, and cell culture treatments. S.W. also generated the final figures for publication. J.R.K. designed and generated the SARS-CoV-2 5′ UTR dual-reporter cell line. F.J.Z. conducted SARS-CoV-2 infection experiments and harvested the infected cells. Q.L. and E.S. performed the mass spectrometry proteomics experiments and deposited the MS data. M.B. conducted mass spectrometry data analysis and figure plotting. J.L. expressed and purified the eIF4A1 protein. N.J.K., M.O., D.L.S., D.R., and K.M.S. acquired funding. D.K., N.J.K., M.O., D.L.S., D.R., K.L., and K.M.S. supervised the research and interpreted data. All authors contributed to editing the manuscript.

---

**Fig. S1.**
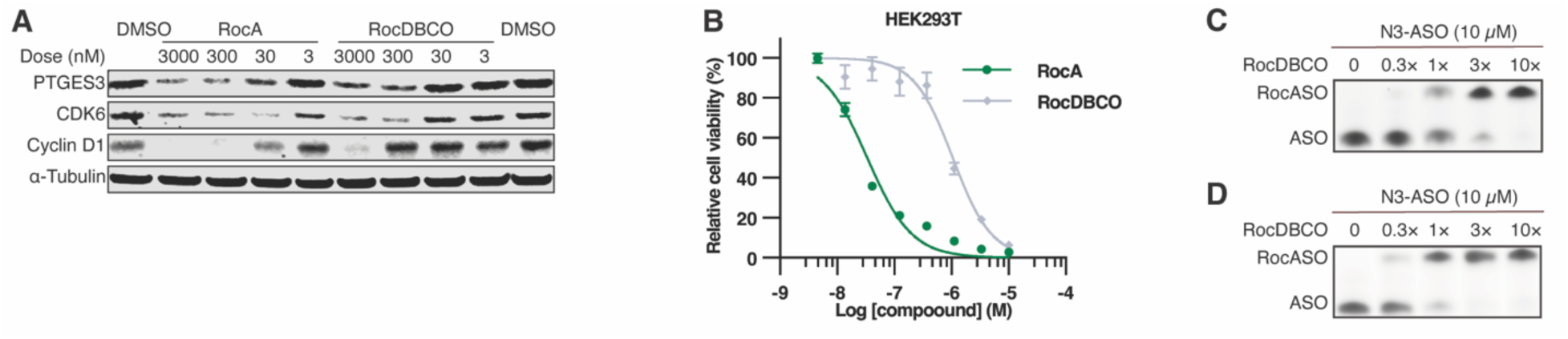
Functional characterization of RocDBCO and its use in RocASO synthesis. **(A)** Immunoblots showing the expression of RocA-sensitive, disease-relevant genes following treatment with RocA or RocDBCO in HEK293T cells for 24 h. **(B)** Cell viability assay of HEK293T cells treated with RocA or RocDBCO for 72 h, data are presented as mean ± SEM (n = 3 biological replicates). All treatments were normalized to 0.1% DMSO control. **(C, D)** Reaction monitoring of the strain-promoted azide-alkyne cycloaddition (SPAAC) reaction between RocDBCO and azide-modified ASO (N3-ASO) by gel electrophoresis. Full consumption of N3-ASO occurred in the presence of 10× excess RocDBCO in 1 hour **(C)** and a 3× excess RocDBCO in 24 hours **(D)**, both at 37°C in nuclease-free water.

**Fig. S2.**
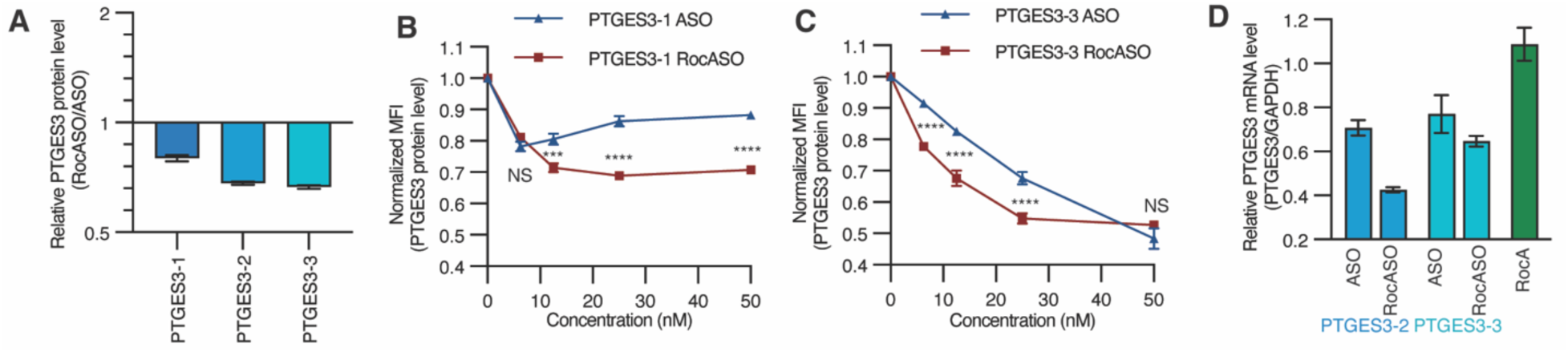
Additional characterization of *PTGES3*-targeting RocASOs. **(A)** PTGES3 fluorescently tagged protein levels after RocASO treatment, normalized to ASO treatment (values <1 indicate enhanced inhibition, 10 nM, 48 hours post-transfection). See Fig. 2D for non-normalized data. **(B, C)** Dose-dependent effects of PTGES3-1 **(B)** and PTGES3-3 **(C)** ASO and RocASO on fluorescently tagged PTGES3 protein levels (48 hours post-transfection). **(D)** *PTGES3* mRNA levels after the indicated treatments (10 nM, 48 hours post-transfection). Unless stated otherwise, data are presented as mean ± SEM (n = 3 biological replicates), and statistical analysis was performed using two-way ANOVA. All treatments are normalized to the vehicle control (transfection reagent + 0.1% DMSO). Flow cytometry data are represented by median fluorescence intensity (MFI).

**Fig S3:**
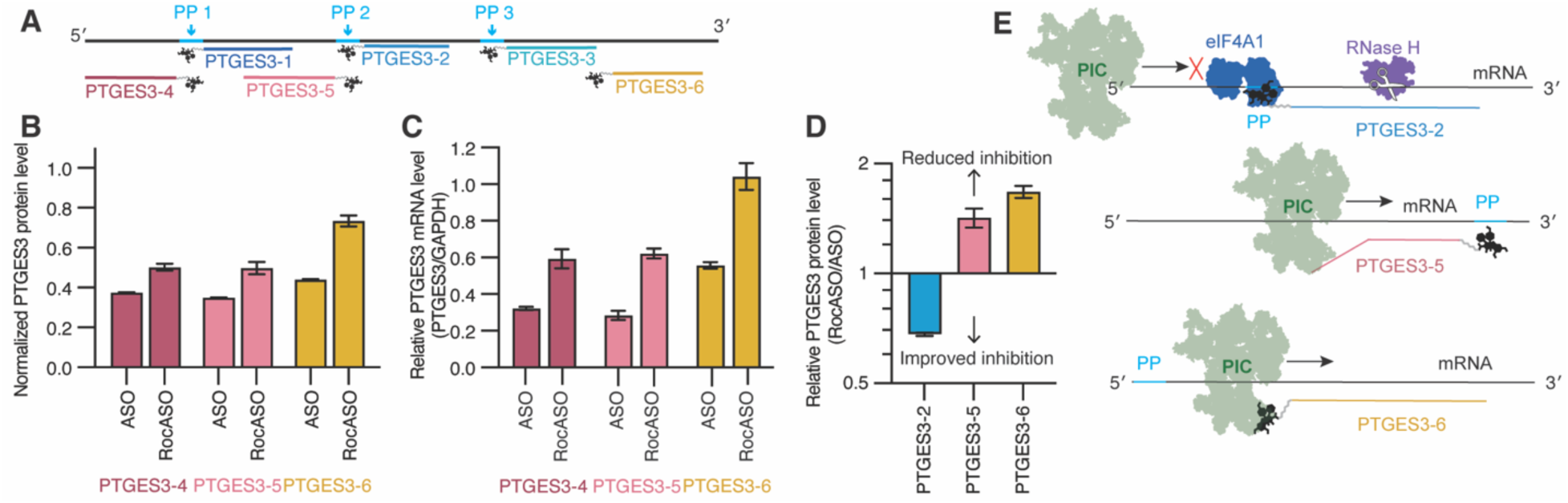
RocA and ASO positioning influences RocASO activity. **(A)** Schematic representation of RocASO design and their targeted sequences relative to defined polypurine motifs (PP, blue). **(B, C)** Non-normalized PTGES3 fluorescently tagged protein levels **(B)** and mRNA levels **(C)** after 48 hours of the indicated treatments (10 nM). **(D)** PTGES3 fluorescently tagged protein levels after RocASO treatment, normalized to ASO treatment (values <1 indicate enhanced inhibition,10 nM, 48 hours post-transfection). See Fig. 2D, fig. S3B for non-normalized data. PTGES3-2 represents a RocASO design with enhanced efficacy, whereas PTGES3-5 and PTGES3-6 represent ineffective designs. **(E)** Schematic illustration of mechanistic models for effective and non-effective RocASO designs. Top panel: RocASOs (PTGES3-2 and -3) with enhanced efficacy clamp eIF4A on PP upstream of the ASO-RNA duplex and block pre-initiation complex (PIC) scanning, leading to effective translation inhibition. Middle panel: RocASOs where ASOs are positioned 5′ to the PP (PTGES3-4, -5) may be displaced by PIC scanning, reducing efficacy. Bottom panel: RocASOs where ASOs are placed too far from a PP (PTGES3-6) prevent RocA from effectively clamping eIF4A. Unless stated otherwise, data are presented as mean ± SEM (n = 3 biological replicates). All treatments are normalized to the vehicle control (transfection reagent + 0.1% DMSO). Flow cytometry data are represented by median fluorescence intensity.

**Fig. S4:**
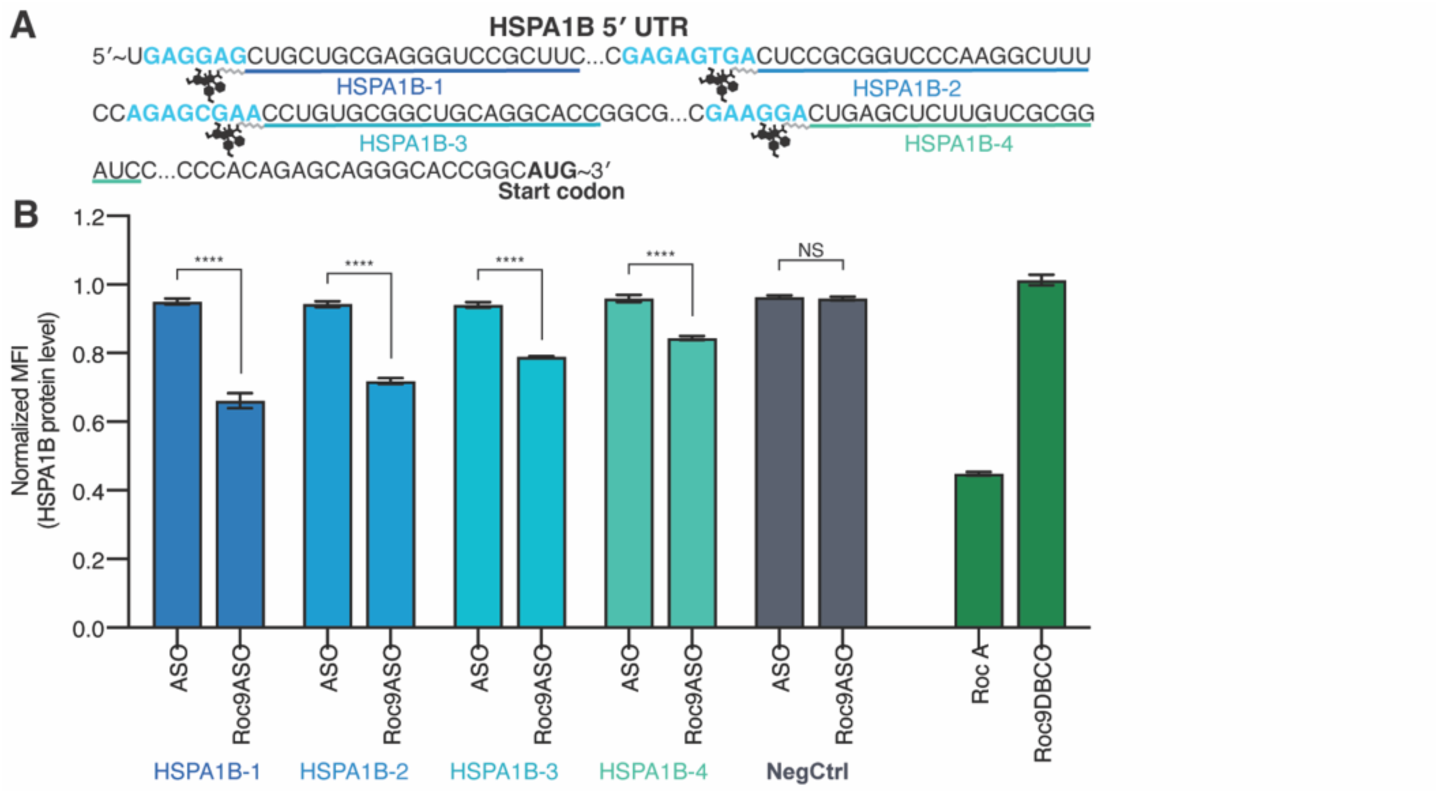
Design and functional characterization of *HSPA1B*-targeting RocASOs. **(A)** *HSPA1B* mRNA 5′ UTR sequence highlighting polypurine motifs (blue) and RocASO target sites (underlined). **(B)** HSPA1B protein expression as measured by fluorescence flow cytometry following indicated treatments (10 nM, 48 hours post-transfection) in HEK293T split-mNeonGreen (mNG) cells, where endogenous HSPA1B has been tagged. Data are presented as mean ± SEM (n = 3 biological replicates), and statistical analysis was performed using two-way ANOVA. All treatments are normalized to the vehicle control (transfection reagent + 0.1% DMSO). Flow cytometry data are represented by median fluorescence intensity (MFI).

**Fig. S5.**
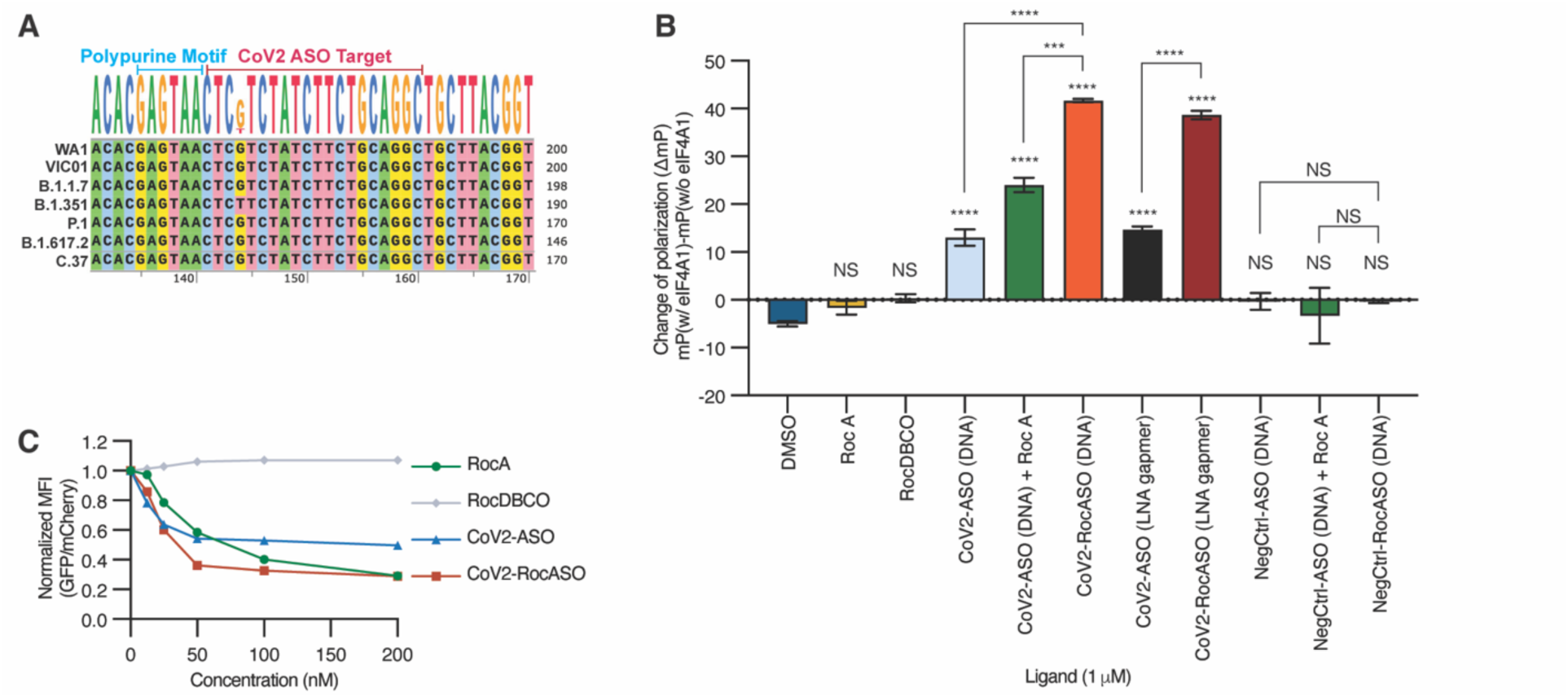
Ternary complex formation with a conserved SARS-CoV-2 target sequence. **(A)** Sequence alignment showing that the CoV2 RocASO targeting region is highly conserved among different SARS-CoV-2 strains. **(B)** Expanded FP assay results from Fig. 3B. Change of polarization values (ΔmP) indicates FP change in the presence or absence of 500 nM eIF4A1 protein with 1 µM of the indicated treatment and 10 mM AMP-PNP. Data are represented as mean ± SEM. **(C)** Dose-dependent effects of RocA, RocDBCO, CoV2-ASO and CoV2-RocASO on the dual-reporter assay system measuring the specific effect of SARS-CoV-2 5′ UTR–driven translation (16 hours post-transfection). All treatments are normalized to the vehicle control (transfection reagent + 0.1% DMSO). Data are presented as mean ± SEM (n = 3 biological replicates), and statistical analysis was performed using two-way ANOVA.

**Fig. S6.**
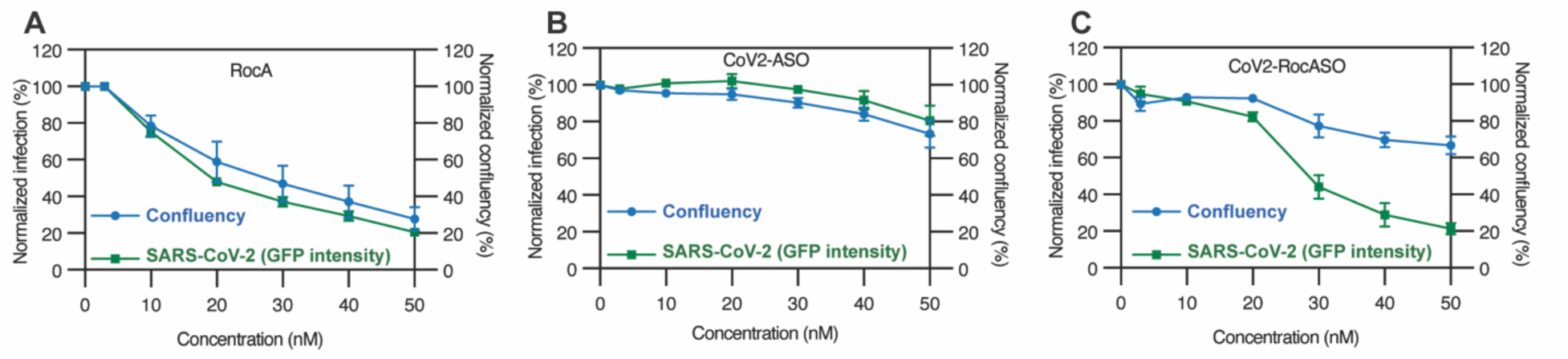
Efficacy and toxicity of different translational inhibition modalities in a cellular model of SARS-CoV-2 infection. Measurements of viral replication and host cell confluency following treatment with RocA **(A)**, CoV2-ASO **(B)**, and CoV2-RocASO **(C)**. All treatments are normalized to the vehicle control (transfection reagent + 0.1% DMSO). Data are presented as mean ± SEM (n = 4 biological replicates), and statistical analysis was performed using two-way ANOVA.

**Fig. S7.**
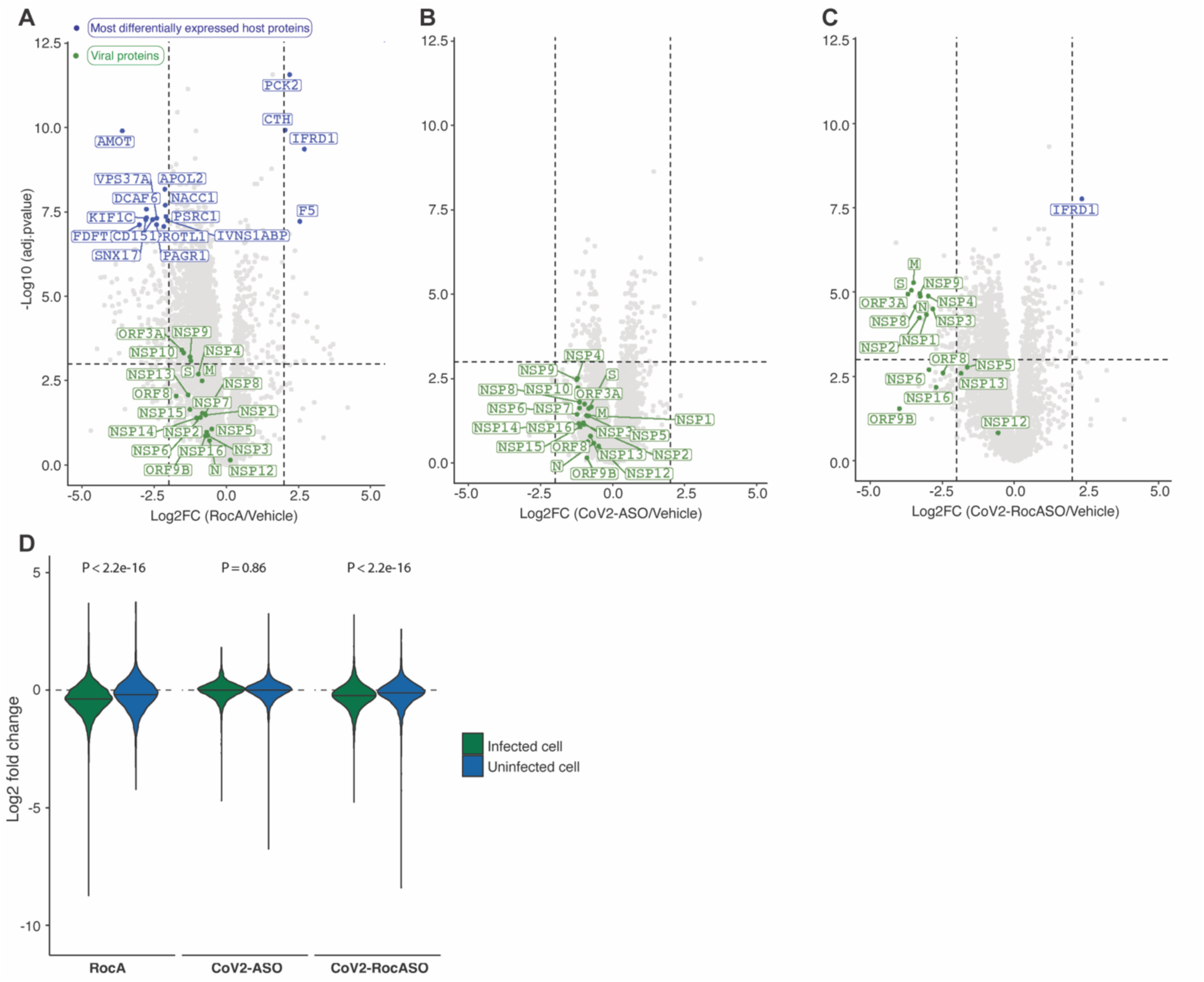
Proteomic analysis of host and viral responses. **(A–C)** Volcano plots showing fold change in host and viral protein expression relative to vehicle control (transfection reagent + 0.1% DMSO) following treatment with RocA **(A)**, CoV2-ASO **(B)**, and CoV2-RocASO **(C)**. Viral proteins are labeled in green; the most significantly differentially expressed host proteins (adj.p < 1e-7, |Log_2_FC| > 2) are labeled in blue. Horizontal and vertical dotted lines indicate significance (adj.p = 0.001) and fold change (log₂FC = ±2) thresholds, respectively (n = 3 biological replicates). **(D)** Violin plots showing the distribution of host protein fold changes under the indicated treatments in either infected or uninfected cells. Horizontal lines represent group medians. Median log₂ fold changes were: RocA infected: –0.43, uninfected: –0.23; CoV2-ASO infected: 0.01, uninfected: 0.02; CoV2-RocASO infected: –0.26, uninfected: –0.11. Statistical comparisons were performed using two-sided t-tests (n = 3 biological replicates).

**Table S1.** Sequences and modifications of oligonucleotides.

| Name | Sequence (5' to 3') |
| --- | --- |
| ASO-Cy5 | Cy5* +G* +G* +C* +T* +G* C* T* G* C* T* A* G* G* G* A*<br>+G* +T* +C* +G* +A* -Azide |
| PTGES3-1 | +C* +T* +T* +T* +T* T* C* T* C* T* C* C* G* G* T* +C*<br>+G* +C* +G* +G* -Azide |
| PTGES3-2<br>(Gapmer) | +G* +G* +C* +T* +G* C* T* G* C* T* A* G* G* G* A* +G*<br>+T* +C* +G* +A* -Azide |
| PTGES3-2<br>(Mixmer) | G* +G* C* +T* G* +C* T* +G* C* +T* A* +G* G* +G* A* +G*<br>T* +C* G* +A* -Azide |
| PTGES3-2<br>(All LNA) | +G* +G* +C* +T* +G* +C* +T* +G* +C* +T* +A* +G* +G*<br>+G* +A* +G* +T* +C* +G* +A* -Azide |
| PTGES3-3 | +C* +G* +G* +G* +C* G* A* A* C* T* G* G* T* G* G* +G*<br>+C* +G* +G* +G* -Azide |
| PTGES3-4 | Azide-* +A* +G* +G* +C* +G* A* C* G* G* C* G* G* C* A*<br>G* +C* +G* +G* +C* +G |
| PTGES3-5 | Azide-* +G* +G* +T* +G* +G* C* G* A* C* T* C* C* G* C*<br>T* +T* +T* +T* +T* +C |
| PTGES3-6 | +G* +G* +C* +A* +G* G* G* G* G* A* C* G* G* G* C* +G*<br>+A* +A* +C* +T* -Azide |
| HSPA1B-1 | +G* +A* +A* +G* +C* G* G* A* C* C* C* T* C* G* C* +A*<br>+G* +C* +A* +G* -Azide |
| HSPA1B-2 | +A* +A* +G* +C* +C* T* T* G* G* G* A* C* C* G* C* +G*<br>+G* +G* +A* +G* -Azide |
| HSPA1B-3 | +G* +G* +T* +G* +C* C* T* G* C* A* G* C* C* G* C* +A*<br>+C* +A* +G* +G* -Azide |
| HSPA1B-4 | +G* +A* +T* +C* +C* G* C* G* A* C* A* A* G* A* G* +C*<br>+T* +C* +A* +G* -Azide |
| CoV2 | +G* +C* +C* +T* +G* C* A* G* A* A* G* A* T* A* G* +A*<br>+C* +G* +A* +G* -Azide |
| CoV2<br>(DNA) | G C C T G C A G A A G A T A G A C G A G -Azide |
| NegCtrl<br>(Fig. 3C, fig. S5B) | C C T A G C C G G C C G C C C G C T C G -Azide |
| NegCtrl<br>(fig. S4B) | +C* +C* +G* +A* +G* C* C* T* G* C* C* G* T* C* C* +G*<br>+C* +C* +C* +G* -Azide |

Cy5: 5' Cy5™ from IDT; \*: The phosphorothioate (PS) bond substitutes a sulfur atom for a non-bridging oxygen in the phosphate backbone; +: Locked nucleic acids; Azide: NHS Ester linked Azide.

## Materials and Methods

### Chemical Synthesis of RocDBCO

Nuclear magnetic resonance (NMR) spectra were recorded on a Bruker spectrometer at 400 MHz. Chemical shifts were reported as parts per million (ppm) from solvent references. Liquid chromatography-mass spectrometry (LC-MS) was performed on a Waters Xevo G2-XS QTof (0.6 mL/min) using an ACQUITY UPLC BEH C18 column (Waters) and a water/acetonitrile gradient (0.05% formic acid) using Optima LC-MS grade solvents (Fisher Scientific). All other solvents were of ACS grade (Fisher Scientific, Millipore Sigma) and used without further purification. Rocagloic acid was obtained from MedChem Express (HY-19355) and DBCO-PEG9-amine was obtained from BroadPharm (BP-24150). Commercially available reagents were used without further purification. Preparative high-performance liquid chromatography (HPLC) was performed on an AutoPurification System using an XBridge BEH C18 OBD Prep Column (Waters) or a CombiFlash EZ Prep using a RediSep C18 Prep HPLC Column (Teledyne ISCO) with a water/acetonitrile gradient.

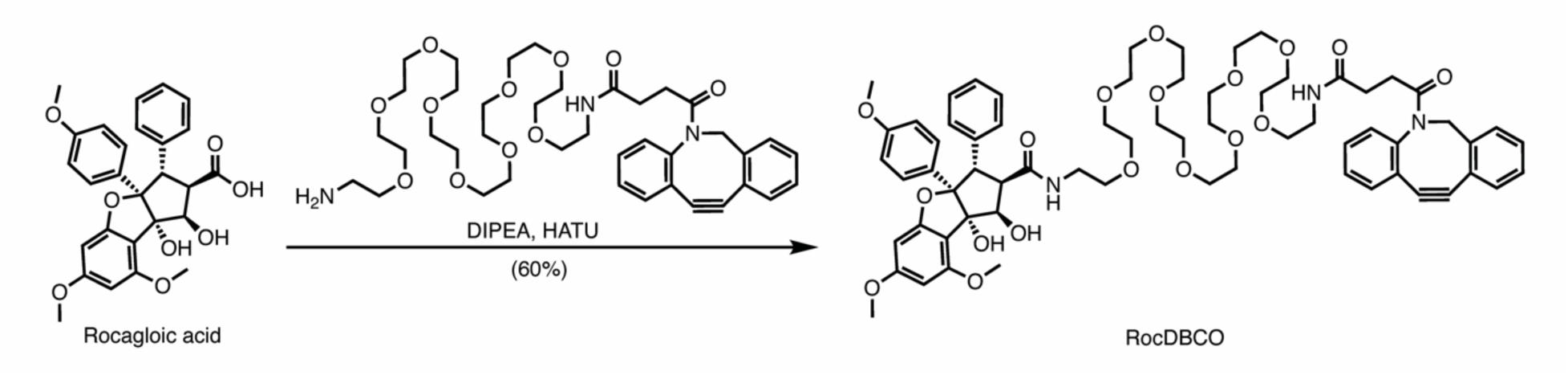

To a mixture of rocagloic acid (10 mg, 0.021 mmol) and DBCO-PEG9-amine TFA salt (17.9 mg, 0.021 mmol) in *N*,*N*-dimethylformamide (0.418 mL) was added *N*,*N*-diisopropylethylamine (15 μL, 0.084 mmol). The solution was cooled in an ice-water bath before the addition of 1-[bis(dimethylamino)methylene]-1*H*-1,2,3-triazolo[4,5-b]pyridinium 3-oxide hexafluorophosphate (7.9 mg, 0.021 mmol) and stirred at room temperature for 1 hour. The crude was purified by reverse-phase HPLC to afford RocDBCO (15 mg, 0.012 mmol, 60% yield) as a white solid.

**^1^H NMR** (400 MHz, DMSO-d_6_) δ 8.29 (t, *J* = 5.6 Hz, 1H), 7.74 (t, *J* = 5.6 Hz, 1H), 7.69 – 7.65 (m, 1H), 7.64 – 7.59 (m, 1H), 7.53 – 7.42 (m, 3H), 7.41 – 7.27 (m, 3H), 7.08 – 6.99 (m, 4H), 6.98 – 6.92 (m, 3H), 6.58 (d, *J* = 9.0 Hz, 2H), 6.26 (d, *J* = 1.9 Hz, 1H), 6.10 (d, *J* = 2.0 Hz, 1H), 5.03 (d, *J* = 14.0 Hz, 1H), 4.94 (s, 1H), 4.60 – 4.50 (m, 2H), 4.17 (d, *J* = 14.1 Hz, 1H), 3.85 (dd, *J* = 14.1, 4.9 Hz, 1H), 3.77 (s, 3H), 3.73 (s, 3H), 3.64 – 3.57 (m, 5H), 3.54 – 3.41 (m, 35H), 3.21 – 3.12 (m, 2H), 3.12 – 3.03 (m, 2H), 2.63 – 2.53 (m, 1H), 2.28 – 2.18 (m, 1H), 2.03 – 1.94 (m, 1H), 1.76 (ddd, *J* = 16.2, 8.0, 5.6 Hz, 1H).

**^13^C NMR** (100 MHz, DMSO-d_6_) δ 171.08, 171.03, 170.27, 162.55, 160.60, 157.88, 157.43, 151.61, 148.43, 138.44, 132.41, 129.61, 128.91, 128.85, 128.70, 128.12, 127.98, 127.92, 127.66, 127.24, 126.78, 125.71, 125.14, 122.53, 121.40, 114.22, 111.69, 108.53, 108.15, 101.27, 93.27, 91.72, 88.40, 78.93, 69.76, 69.71, 69.68, 69.59, 69.54, 69.06, 68.99, 55.44, 55.35, 55.26, 54.88, 54.74, 49.99, 30.33, 29.70.

**HRMS** (*m/z*): calculated for C66H82N3O18+ [M + H]+ 1204.5588, found 1204.5645.

### RocASO Conjugation

#### Click Chemistry Conjugation

RocDBCO and azide-modified antisense oligonucleotides (ASOs) were conjugated using strain-promoted azide–alkyne cycloaddition (SPAAC) click reactions. Reaction mixtures were prepared by combining ASO and RocDBCO at specified molar ratios. Unless otherwise indicated, reactions were performed at a RocDBCO:ASO molar ratio of 10:1 in a final volume of 50 µL, adjusted with RNase-free water. Reactions were incubated at 37°C for specified durations (default: 1 hour), followed by purification of the resulting RocASO conjugates using an Oligo Clean & Concentrator kit (D4061, Zymo Research) according to the manufacturer’s protocol. The absorbance at 260 nm (A260) of purified conjugates was measured using a NanoPhotometer NP80 spectrophotometer (Implen) to determine concentrations. Products were used immediately or stored at -20°C for further analysis.

#### Gel Electrophoresis Analysis

Conjugation efficiency was verified by gel electrophoresis. Briefly, aliquots containing 1.5 nmol of purified conjugate were mixed with gel-loading dye (B7024S, New England Biolabs) and RNase-free water to a final volume of 12 µL. Samples (10 µL each) were resolved on 15% TBE-Urea gels (EC68852BOX, Thermo Fisher Scientific) in 1× TBE buffer at 100 V for 90 min. Gels were stained for 30 min in the dark with SYBR™ Gold nucleic acid stain (S11494, Fisher Scientific; diluted 1:10,000 in 1× TBE buffer). Conjugation products were visualized under blue-light illumination.

### Cell culture and reagents

HEK293T cells were a generous gift from the laboratory of Luke Gilbert (University of California, San Francisco). HEK293T-ACE2/TMPRSS2 (HEK293T-AT) cells were kindly provided by Jennifer Doudna’s laboratory (Gladstone Institute). The split mNG endogenously tagged HEK293T cell lines were obtained from Manuel D. Leonetti’s laboratory (Chan Zuckerberg Biohub). All cell lines were cultured in DMEM high-glucose medium (11995073, Gibco) supplemented with 10% (v/v) fetal bovine serum (F-0500-D, Atlas Biologicals), and 1% (v/v) penicillin-streptomycin (15070063, Gibco). Cells were maintained at 37°C in a humidified incubator with 5% CO₂ and routinely tested to ensure the absence of mycoplasma contamination.

Rocaglamide was obtained from MedChem Express (HY-19356). All the ASOs were purchased from Integrated DNA Technologies. Small molecules were stored at -20°C as 10 mM stock solutions in dimethyl sulfoxide (DMSO). ASOs were stored at -20°C as 100 µM stock solutions in Ultrapure DNase/RNase Free Distilled Water (10977015, Fisher Scientific).

### Protein expression and purification

The DNA sequence encoding human eIF4A1 was codon-optimized for protein expression in E. coli and synthesized by Genscript. It was then cloned into the pET21a(+) vector with an N-terminal TEV-cleavable 6×His tag.

Chemically competent BL21(DE3) cells (C2527H, NEB) were transformed with the plasmid pET21a(+)-His-TEV-eIF4A1 and plated onto LB agar plates containing 100 µg/mL ampicillin, then incubated overnight at 37°C. A single colony was inoculated into Terrific Broth supplemented with 100 µg/mL ampicillin and cultured at 37°C with shaking at 200 rpm until the optical density at 600 nm (OD_600_) reached approximately 0.6. Protein expression was induced with 0.5 mM IPTG at 18°C overnight. Cells were harvested by centrifugation at 4,500 × g for 15 min, resuspended in lysis buffer (20 mM HEPES, pH 7.5, 500 mM NaCl, 10 mM imidazole, 2 mM DTT), and lysed by sonication. The lysate was clarified by high-speed centrifugation at 19,000 × g for 60 min. The clarified supernatant was subjected to immobilized metal affinity chromatography using Co-TALON resin (Clontech, Takara Bio USA; 4 mL slurry per liter of culture) and incubated for 1 hour at 4°C with gentle end-to-end mixing. After incubation, the resin was washed three times, each time with 10 mL of lysis buffer. His-tagged eIF4A1 was eluted with 15 mL of elution buffer (20 mM HEPES, pH 7.5, 500 mM NaCl, 250 mM imidazole, 2 mM DTT). The N-terminal His-tag was cleaved using His-tagged TEV protease (Macrolab, UC Berkeley) under dialysis conditions (20 mM HEPES, pH 7.5, 200 mM NaCl, 2 mM DTT) overnight at 4°C. After cleavage, the mixture was incubated with HisPur Ni-NTA resin (Thermo Fisher Scientific; 1 mL slurry per liter of culture) for 10 min, and the flow-through containing tag-free eIF4A1 was collected. The collected protein solution was concentrated to approximately 20 mg/mL using a centrifugal concentrator with a 10 kDa molecular weight cut-oi (Vivaspin Turbo 15, Sartorius), then further purified by size exclusion chromatography on a Superdex 200 10/300 GL column (GE Healthcare Life Sciences) using SEC buffer (20 mM HEPES, pH 7.5, 150 mM NaCl, 1 mM DTT). Pure fractions containing eIF4A1 were pooled, concentrated to around 5 mg/mL, and stored at -80°C. This purification protocol typically yields 5–10 mg of purified eIF4A1 protein per liter of bacterial culture. The molecular weight of the purified eIF4A1 protein was confirmed by LC-MS to be 46,081 Da, consistent with the calculated molecular weight of 46,080 Da.

### Fluorescent polarization assay

The RNA probe used in this assay consisted of a 37-nucleotide sequence derived from SARS-CoV-2 mRNA, encompassing both the antisense oligonucleotide (ASO) binding region and the adjacent polypurine sequence. The probe was labeled at its 5′ terminus with fluorescein amidite (FAM). The RNA probe sequence is as follows: 5′ -FAM-UGA CAG GAC ACG AGU AAC UCG UCU AUC UUC UGC AGG C - 3′.

For each reaction, 10 nM of the 5′-FAM-labeled RNA probe was incubated with the indicated ligand (1 µM final concentration) in assay buffer containing 20 mM HEPES (pH 7.5), 150 mM NaCl, 1 mM DTT, 1 mM AMP-PNP, and 1 mM MgCl₂. Assays were performed in a final volume of 10 µL in the presence or absence of 500 nM recombinant eIF4A1 protein. Reaction mixtures were incubated at room temperature for 60 min, after which fluorescence polarization was measured using a Spark multimode plate reader (Tecan) at room temperature.

### Plasmids and viral transduction

The fluorescent translational reporter has been previously published (*37*). The SARS-CoV2 sequence was comprised of the 5′ UTR plus the first 30bp of Orf1a to maintain the RNA structure and was purchased as a gBlock (IDT). The sequence was based on the USA-WA1/2020 strain. The SARS-CoV-2 sequence and destabilized GFP were cloned into the MluI site using multi-piece In-Fusion cloning to ensure seamless insertion. The destabilized GFP sequence was cloned from FUGW-d2GFP-ZEB1 3’UTR (79601, Addgene). All cloning and plasmids were transformed into either lab made DH5a or Stellar (Takara) competent bacteria.

To identify potential polypurine motifs within the SARS-CoV-2 sequence, we used FIMO within the MEME suite (*38*) to search for 6-mer motifs as identified by eIF4A1 RNA interaction studies (*16*).

To generate the translational reporter virus, HEK293T were transfected with 9μg lentiviral expression construct, 7 μg psPAX2, and 2.5 μg PMD2.G. Transfections were done in 10 cm plates using Lipofectamine 2000 (Life Technologies). Viral supernatant was collected at 48 and 72 hours after transfection and concentrated using Lenti-X Concentrator (Takara). To generate the reporter cell lines, HEK293T cells were seeded into lentiviral-containing media and transduced overnight with 5 μg/mL polybrene with fresh media change the next morning. Cells were selected using 5 μg/mL blasticidin approximately 36 hours after initial infection. To obtain a cell population with robust reporter expression (high GFP and high mCherry fluorescence), the cells were sorted using a BD Aria 3 sorter.

### Transfection

Cells were transfected using Lipofectamine™ RNAiMAX (13-778-150, Thermo Fisher Scientific) in a 24-well plate format. One day prior to transfection, cells were seeded at 50,000 cells per well in 500 µL of antibiotic-free growth medium to achieve approximately 30–50% confluency at transfection.

On the day of transfection, ligands and Lipofectamine™ RNAiMAX (1 µL/well) were diluted in 50 µL of Opti-MEM® I Reduced Serum Medium (31-985-062, Thermo Fisher Scientific) separately and mixed gently. Both solutions were combined, mixed gently, and incubated for 15 minutes at room temperature to allow complex formation. 100 µL of RNAi-Lipofectamine complexes were then added to each well, resulting in a total medium volume of 600 µL per well. Cells were incubated at 37°C with 5% CO₂ for the indicated duration before analysis of gene knockdown.

For transfection in other plate formats, the amounts of Lipofectamine™ RNAiMAX, cells, and medium were scaled proportionally based on surface area.

### Fluorescence reporter assay

Following the indicated treatment duration, cells were harvested by trypsinization, and the reaction was neutralized by the addition of growth medium. Cells were then centrifuged at 300 × *g* for 5 minutes and washed twice with ice-cold phosphate-buffered saline (PBS). If required, cell pellets were divided for parallel protein and RNA analyses. For flow cytometry analysis, cells were resuspended in ice-cold fluorescence-activated cell sorting (FACS) buffer (PBS containing 1% BSA and 0.05% sodium azide) supplemented with 1 µM SYTOX™ Blue dead cell stain (S34857, Thermo Fisher Scientific). Fluorescence was assessed immediately on a CytoFLEX flow cytometer (Beckman Coulter). Data analysis was performed using FlowJo software.

### qRT-PCR analysis

RNA from treated cells was isolated using the RNeasy Plus Mini Kit (QIAGEN) according to manufacturer instructions. Reverse transcription was performed using the SuperScript III First-Strand Synthesis SuperMix for qRT-PCR (Life Technologies) according to manufacturer instructions. The reverse transcription products were evaluated by qRT-PCR using the Maxima SYBR Green qPCR Master Mix (Life Technologies) on a Bio-Rad CFX Touch Real-Time PCR system according to manufacturer instructions. GAPDH served as a reference gene. All samples were evaluated using the ΔΔCq method under the gene expression tab in the Bio-Rad CFX Maestro for Mac 1.1 software. Primer sequences are as follows: GAPDH forward: 5′-AAT CCC ATC ACC ATC TTC CA-3′; GAPDH reverse: 5′-TGG ACT CCA CGA CGT ACT CA-3′; PTGES3 forward: 5′-GAT TCC AAG CAT AAA AGA ACG GAC-3′; PTGES3 reverse: 5′-GAA TCA TCT TCC CAG TCT TTC CAA-3′.

### Immunoblotting

Cells were seeded into 6-well plates and incubated overnight at 37°C. Following treatment with compounds at the indicated concentrations and durations, cells were placed on ice and washed twice with PBS. Unless otherwise indicated, the cells were scraped with a spatula, pelleted by centrifugation (500 × g for 5 min), and lysed in lysis buffer [100 mM Hepes (pH 7.5), 150 mM NaCl, and 0.1% NP-40] supplemented with cOmplete Protease Inhibitor Cocktail Tablets (Roche) and PhosSTOP (Roche) on ice for 10 min. Lysates were clarified by high-speed centrifugation (22,000 × g for 10 min). Concentrations of lysates were determined with a protein BCA assay (Thermo Fisher Scientific) and adjusted to the same concentration with additional RIPA buffer. Samples were mixed with 5× SDS loading dye and heated at 95°C for 5 min. SDS–PAGE was run with Novex 4–12% Bis-Tris gel (Invitrogen) in MES running buffer (Invitrogen) at 140 V for 90 min following the manufacturer’s instructions. Protein bands were transferred onto nitrocellulose membranes by iBlot 3 Western Blot Transfer System (Thermo Fisher Scientific) and blocked using blocking buffer [5% bovine serum albumin (Millipore) in Tris-buffered saline, 0.1% Tween 20 (TBST) supplemented with 0.02% NaN3]. Membranes were probed with primary antibodies against PTGES3 (HPA038672) from Sigma-Aldrich, CDK6 (3136S), Cyclin D1 (2978S), and Tubulin (3873S) from Cell Signaling Technology diluted (1:1000) in blocking buffer. After primary antibody incubation, membranes were treated with IRDye secondary antibodies (926-68070 or 926-32211) from LI-COR Biosciences according to manufacturer’s recommendations and scanned on an ChemiDoc MP (Bio-Rad).

### Cell viability assays

Cells were seeded into white, flat–bottom, opaque 96-well plates and incubated at 37°C overnight. Cells were treated with the indicated concentrations of compound in 9-point 3-fold dilution series (100 μL final volume per well) and incubated at 37°C for 72 hours. Cell viability was assessed by CellTiter-Glo (CTG) Luminescent Cell Viability Assay (Promega). Cells were equilibrated to room temperature before the addition of CTG reagent (100 μL per well). Plates were agitated on an orbital shaker and luminescence signal was measured on a Spark (Tecan) plate reader. Repeated measurements of luminescence were performed as technical replicates to determine incubation times optimal for signal-to-noise.

Luminescence measurements were normalized to DMSO-treated controls to determine relative cell viability.

### SARS-CoV-2 *in vitro* antiviral assay

A fluorescent SARS-CoV-2 infectious clone (ic-SARS-CoV-2-mNG), containing an mNeonGreen reporter instead of Orf7a/b (*39*), was propagated in Vero-TMPRSS2 cells and subsequently used for antiviral screening assays performed with the Incucyte live-cell imaging system. One day prior to treatment, the HEK293T-AT cells were seeded onto poly-L-lysine coated 96-well plates. ASO and RocASO transfections were performed as described above, and cells were incubated for 24 hours. The following day, cells were infected with ic-SARS-CoV-2-mNG at a multiplicity of infection (MOI) of 0.1 for 2 hours at 37 °C and 5% CO_2_ under BSL-3 conditions. Antiviral activity was assessed using the Incucyte system as previously described (*40*). Infected cells were imaged every hour for 24 hours using a 10× objective. Three images per well were acquired under standard cell culture conditions (37°C, 5% CO_2_). Viral infection was quantified by measuring total green object integrated intensity (GCU × μm²/image) with a 300 ms exposure time. Cell growth was assessed using phase-contrast imaging. Phase analysis used AI-based confluence segmentation. Green-channel analysis was performed using Top-Hat segmentation (50 μm radius, 0.5 GCU threshold, edge split enabled). Red-channel analysis used similar parameters, with a 100 μm radius and a 1 RCU threshold. Antiviral efficacy was calculated as percent infection normalized to vehicle control. Cell growth was expressed as percent confluency normalized to the same controls. Specific antiviral activity was determined by calculating the infection/confluency ratio, normalized to vehicle control. A previously validated ASO was included as a positive control in all assays (*41*). Experiments were performed in triplicate with four biological replicates per condition.

### MS sample preparation

Cell pellets were resuspended in lysis buffer composed of 150 mM sodium chloride, 1.0% NP-40, 0.5% sodium deoxycholate, 0.1% sodium dodecyl sulfate, 50 mM Tris, pH 8.0. Lysate protein concentration was measured by Bradford Assay. The reduction of protein disulfide bonds was performed by the addition of DTT to a final concentration of 5 mM, and incubation at room temperature for 30 min. Next, cysteine residues were alkylated by the addition of iodoacetamide to a final concentration of 10 mM for 30 min at room temperature in the dark. The alkylation reaction was quenched by the addition of DTT to a final concentration of 5 mM and incubated for 30 min at room temperature. Buffer exchange and proteolytic digestion of each sample was performed on a KingFisher Flex unit using a protein aggregation capture approach. Here, the 96-well comb is stored in plate #1, magnetic carboxylated beads and the lysate in plate #2, 95% acetonitrile wash solutions in plates #3 to #5, 70% ethanol wash solutions in plates #6 and #7, elution/digestion buffer with the digestion enzymes in plate #8. A volume of 100 µL of lysate (50 µg of protein) was added to plate #2, followed by the addition of 20 µL of MagReSyn Amine beads (Resyn Biosciences), and lastly 280 µL of acetonitrile. Protein aggregation onto the beads was then induced over the course of 24 min by 4 cycles of mixing (1 min), pause (5 min), and collection of the magnetic beads. Next, beads with bound protein were transferred to plate #3 and released into 150 µL of 95% acetonitrile and washed by 5 cycles of mixing. This process was repeated twice more in plates #4 and #5. Bounds beads were then transferred to plate #6 and released into 150 µL of 70% acetonitrile and washed by 5 cycles of mixing. This process was repeated once more in plate #7. Lastly, the beads containing the bound protein were released into 150 µL of 50 mM ammonium bicarbonate (pH 7.8) containing both Lys-C and trypsin (1:100 enzyme: protein ratio for each). The plate was then sealed, and proteolytic digestion proceeded overnight (18 hours) at 37 °C with agitation on an Eppendorf ThermoMixer at 800 rpm. Lastly, the resulting peptides were filtered through 0.45 µm membranes and dried by vacuum centrifugation.

### MS data acquisition and analysis

For MS analysis, peptide samples were resuspended with 100 µL of 0.1% formic acid (FA) to a concentration of 0.5 µg/µL and 1 µL of was injected for analysis into an Orbitrap Exploris 480 mass spectrometer coupled with a Vanquish Neo liquid chromatography system. Peptides were separated on a PepSep column (15 cm x 150 µm x 1.5 µm) using an 80 min separation gradient at a flow rate of 500 nL/min, with a gradient from 3% to 30% mobile phase B over 60 min, followed by an increase to 40% B over 10 min, and held at 95% B for 10 min. Mobile phase A was 0.1% FA, while mobile phase B was 0.1% formic acid in 80% acetonitrile. Data-independent acquisition (DIA) mode was used for MS analysis with the following parameters. The precursor’s full scan was performed in the Orbitrap at the resolution of 60,000 over the range of 350-950 *m*/*z*, with an AGC target of 300% and an automatic maximum injection time. Following the MS full scan, DIA MS/MS scan was performed with a 15,000 scan resolution, standard AGC target, automatic maximum injection time, and using a normalized HCD collision energy of 28%. MS/MS scans were collected over the range of 350-950 *m*/*z*, with an isolation window of 20 *m*/*z* and a 0.5 *m*/*z* overlap.

The DIA raw data were analyzed with Spectronaut (version 19.4.241120.62635) in a direct-DIA mode against the SwissProt Human database (downloaded 12/12/2024), with SARS-CoV-2 proteins included. The default settings in Spectronaut were used, with fixed modification as cysteine carbamidomethylation, and variable modification as methionine oxidation and protein N-termini acetylation. To keep the real peptide intensities, the imputation and cross-normalization in the default settings were turned oi. Label-free quantification of peptide data, including summarization of peptide intensities to protein intensities and differential expression analysis, was performed using the MSstats R package (v4.14.1). Peptide data was first normalized by equalizing median peptide intensity across all mass spectrometry runs. Peptides only identified in a single run were deemed uninformative and removed before protein summarization and no imputation of missing data was performed. Outlier peptides were excluded from the Tukey median polish calculation to obtain protein intensities from peptide intensities. Pairwise differential expression analysis was done with MSstats to compare all treatment and infection conditions, selected comparisons are shown. The differential p-values were adjusted using Benjamini & Hochberg (BH) correction. The differential protein set for each comparison is defined as those proteins having adj.p < 0.05 and absolute log_2_FC > 1.

The viral protein abundance summary shown in Fig 4C was calculated as the mean of absolute viral protein intensities within each replicate, while the percent of viral protein abundance was calculated for each viral protein (each row) as a percentage of the replicate with the highest viral protein intensity. The density plots in Fig. 4D–F and the violin plots in fig. S7D show the log_2_FCs of treatment effect only for proteins with no replicates missing in the greater condition of the contrast. The following packages in R (v4.4.2, release ‘Pile of Leaves’) were used for figure generation: ggplot2 (v3.5.1), ggrepel (v0.9.6), ComplexHeatmap (v2.22.0), viridis (v0.6.5).

The mass spectrometry proteomics data have been deposited to the ProteomeXchange Consortium via the PRIDE (*42*) partner repository with the dataset identifier PXD065291.

